# miRTarDS: High-Accuracy Refining Protein-level MicroRNA Target Interactions from Prediction Databases Using Sentence-BERT

**DOI:** 10.1101/2024.05.17.594604

**Authors:** Baiming Chen

**Affiliations:** School of Medicine, The Chinese University of Hong Kong, Shenzhen, Longgang District, Shenzhen 518172, China; Shenzhen X-Institute, Nanshan District, Shenzhen, Guangdong Province 518055, China

**Keywords:** MicroRNA (miRNA), Sentence-BERT, MicroRNA Target Interactions (MTIs), Gene Regulation, Database

## Abstract

MicroRNAs (miRNAs) regulate gene expression by binding to mRNAs, inhibiting translation, or promoting mRNA degradation. miRNAs are of great importance in the development of various diseases. Currently, numerous sequence-based miRNA target prediction tools are available, however, only 1% of their predictions have been experimentally validated. In this study, we propose a novel approach that leverages disease similarity degree between miRNAs and genes as a key feature to further refine human sequence-based predicted miRNA target interactions (MTIs). To quantify the similarity degree of diseases, we fine-tuned the Sentence-BERT model. Our method achieved an F1 score of 0.88 in accurately distinguishing human protein-level experimentally validated MTIs (functional MTIs, validated through western blot or reporter assay) and predicted MTIs. Moreover, this method exhibits exceptional generalizability across different databases. We applied the proposed method to analyze 1,220,904 human MTIs sourced from miRTarbase, miRDB, and miRWalk, encompassing 6,085 genes and 1,261 pre-miRNAs. Our model was trained in miRTarBase 2022. However, we accurately identified 90% (518/574) of the updated functional MTIs in miRTarbase 2025. This study has the potential to provide valuable insights into the understanding of miRNA-gene regulatory networks and to promote advancements in disease diagnosis, treatment, and drug development.

## 1 Introduction

In 1993, miRNA (Lin-4) was first discovered and found to be critical for the timing of embryonic development in C. elegans [1]. Since then, a large number of miRNAs have been discovered with the development of high-throughput bioinformatics technologies [2]. miRNAs are classified as small non-coding RNAs (ncRNAs), and play a crucial role in the regulation of various physiological processes, such as cell cycle [3], cell growth, development, differentiation, and apoptosis [4], as well as pathological processes [5]. In addition, miRNAs may be promising biomarker candidates for early detection or prognosis of various diseases [6]. Therefore, the identification of MTIs holds biological significance.

There are many existing MTI prediction databases. miRWalk identifies consecutive complementary subsequences between miRNAs and genes to construct a database for MTIs prediction in different species [7]. miRDB considers miRNA over-expression and CLIP-seq data, utilizing a support vector machine (SVM) for MTI prediction in different species [8]. mirSVR trains a regression model to rank miRNA target sites based on the down-regulation score, using data from miRanda [9]. Deep-Target uses Recurrent Neural Networks (RNN) for learning sequence-sequence interaction patterns to predict MTIs [10]. TargetScan predicts MTIs based on seed match and conservativeness analysis in different species [11]. The RNA22 is designed for the identification of miRNA binding sites and their corresponding miRNA and mRNA complexes [12]. PITA predicts miRNA targets by considering both the binding free energy and the accessibility of the target site within the mRNA secondary structure [13]. StarBase explores miRNA-mRNA and miRNA-lncRNA interaction networks based on CLIP-Seq data [14].

However, limited research has focused on assessing the degree of disease similarity between miRNAs and genes. If a specific miRNA and gene are associated with the same or closely related diseases, it is higher possibility that the miRNA and the gene may encounter within the human body. Furthermore, if sequence-based MTI prediction databases suggest a potential interaction between them, it can be inferred that the miRNA is highly likely to silence the gene. There are many miRNA [15] and gene [16] associated disease databases that contain a lot of semantic disease information. So, we use NLP method to explore the degree of disease similarity between miRNAs and genes. For details, we use the Sentence-BERT (SBERT) model to calculate disease similarity. SBERT improves the pre-trained BERT model by incorporating Siamese and triplet network structures [17]. BERT, a bidirectional transformer model, captures the semantic meaning of each word based on its preceding and following context. Traditional BERT requires sentence pairs to be concatenated and processed together, making tasks like sentence similarity calculations computationally expensive. SBERT, by contrast, generates independent sentence embeddings that can be precomputed, allowing for sentence comparisons through simple similarity calculations such as cosine similarity. This approach significantly enhances inference speed. SBERT outperforms both BERT [18] and RoBERTa [19] in terms of speed while maintaining comparable accuracy.

The present study proposes a method for calculating the similarities between miRNAs associated diseases and genes associated diseases, aiming to further refine sequence-based MTI prediction. We assume that protein-level experimentally validated MTIs (functional MTIs, validated through western blot or reporter assay) exhibit different disease similarity patterns from predicted MTIs. The experimental results confirm this hypothesis. The miRTarDS database can be accessed through Github.

## 2 Materials and Methods

### 2.1 Overview

We present a flow chart (Figure 1) outlining the major steps of our study. We first integrated MTI data from miRTarBase, miRWalk, and miRDB. To fine-tune the SBERT model, we constructed the MeSHDS dataset. For each MTI, we computed a disease similarity matrix by inputting the diseases associated with the miRNAs and genes. We then analyzed the distribution of similarity values within the matrix and randomly selected an equal number of protein-level interaction and prediction interaction MTIs to train a random forest classifier. Finally, we applied the classifier to all collected MTIs.

**Figure 1.**
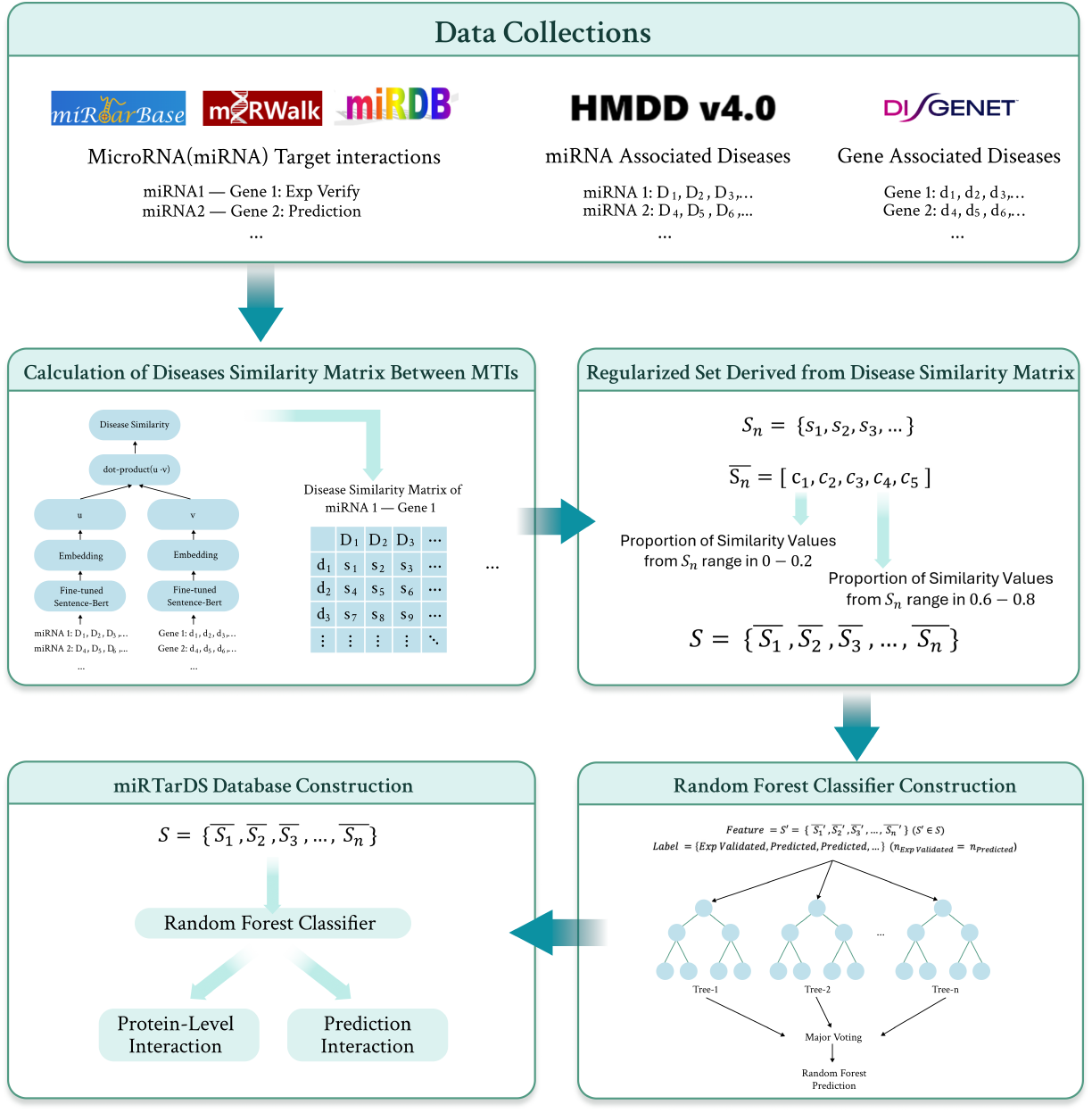
The flow chart illustrates the methodology of the study. Here, *D*_*n*_ de-notes the miRNA-associated disease, *d*_*n*_ represents the gene-associated disease, and *s*_*n*_ denotes the similarity value calculated between *D*_*n*_ and *d*_*n*_. The regularized set of *S*_*n*_, denoted as 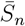, is obtained by transforming the similarity values into discrete intervals. Additionally, *S* represents a comprehensive set containing disease similarity information for all MTIs. From *S*, an equal number of experimentally validated MTIs (*Exp V alidated*) and predicted MTIs (*Predicted*) are randomly sampled to form a subset *S*^*′*^, which is used for meachine learning model construction.

### 2.2 Data Collection

To fine-tune the SBERT model, we downloaded the MeSH (Medical Subject Headings) descriptors (https://meshb.nlm.nih.gov/) from the National Library of Medicine. The MeSH descriptors are standardized terms utilized for indexing and retrieving biomedical and health-related information, forming a hierarchical vocabulary tree (Figure 3). To test the SBERT model performance, we obtained the BIOSSES dataset (Figure 2), which consists of biomedical statements with manually annotated text similarity scores provided by experts [20]. It can be used to evaluate the biomedical text similarity calculation model. We utilized the miR-TarBase [21], miRWalk [7] and miRDB [8] to obtain human protein-level experimentally validated MTIs and predicted MTIs. miRTarBase provides experimentally validated MTIs, which are primarily categorized into two types: Functional MTIs and Functional MTIs (Weak). Functional MTIs are validated through reporter assays, western blot. Functional MTIs (Weak) are identified by microarray, qPCR, or sequencing. We use MTIs in the miRTarBase 2022 as a part of the training set and use updated MTIs in miRTarBase 2025 as a validation set. miRWalk and miRDB provide sequenced-based predicted MTIs. For acquiring information on miRNA-associated and gene-associated diseases, we employed HMDD [15], DisGeNET [16] and KEGG [22] databases. The HMDD provides data on diseases associated with pre-miRNAs. Therefore, for disease associations involving mature miRNAs such as hsa-miR-122b-3p and hsa-miR-122b-5p, we will refer to their corresponding pre-miRNA, hsa-mir-122b, for subsequent calculations. The DisGeNET and the KEGG databases provide diseases associated with genes.

**Figure 2.**
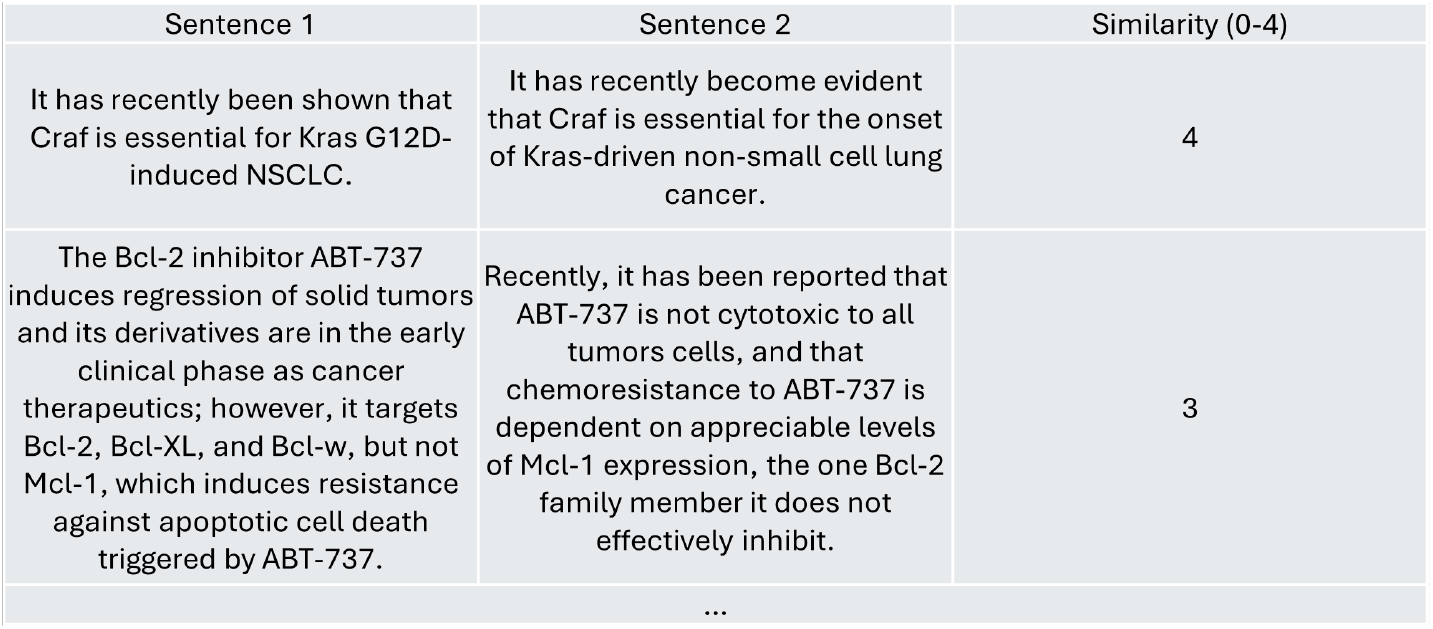
Example of BIOSSES Dataset. BIOSSES dataset is a benchmark dataset specifically designed for evaluating semantic similarity in biomedical texts. The dataset consists of 100 sentence pairs extracted from biomedical literature, each independently scored by five domain experts based on their semantic similarity.

**Figure 3.**
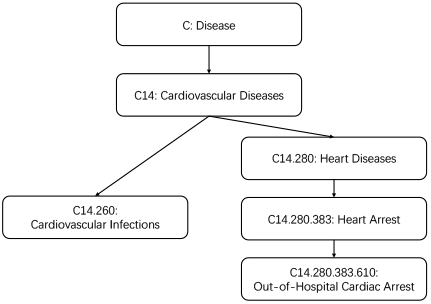
Disease Descriptor ‘Car-diovascular Infections’ and ‘Out-of-Hospital Cardiac Arrest’ Hierarchical Structure in MeSH Tree

### 2.3 Model Construction

#### 2.3.1 Selection of Pretrained Model

The ideal model we seek must fulfill two primary criteria: exceptional performance in semantic similarity calculation tasks and rapid execution speed to facilitate large-scale data processing (Table 1). Moreover, scalability for handling lengthy biomedical semantic strings is favorable. Based on the SBERT website (https://www.sbert.net/), we have selected the ‘multi-qa-MiniLM-L6-cos-v1’ [23] model for further investigation. This model excels in execution speed, and robust semantic similarity calculations, and supports extended input sequences, with a maximum sequence length of 512 compared to 256 for the ‘all-MiniLM-L6-v2’ model.

**Table 1:** Performance Comparison of Pretrained SBERT Models. The **Rank** column indicates the ranking of the models based on their average performance in sentence embedding and semantic search tasks. The **Emb. Perf** column represents the average performance of the model in embedding sentences across different domains, where higher values indicate better performance. The **Sem. Search Perf**. column shows the average performance of the model in semantic search tasks, with higher values indicating better performance. The **Enc. Spd**. column measures the encoding speed of the models in sentences per second on a V100 GPU.

| **Rank<sup>a</sup>** | **Model Name** | **Emb. Perf.<sup>b</sup>** | **Sem. Search Perf.<sup>c</sup>** | **Enc. Spd.<sup>d</sup>** |
| --- | --- | --- | --- | --- |
| 1 | all-mpnet-base-v2 | 69.57 | 57.02 | 2800 |
| 3 | all-distilroberta-v1 | 68.73 | 50.94 | 4000 |
| 4 | all-MiniLM-L12-v2 | 68.70 | 50.82 | 7500 |
| 6 | all-MiniLM-L6-v2 | 68.06 | 49.54 | 14200 |
| 7 | multi-qa-MiniLM-L6-cos-v1 | 64.33 | 51.83 | 14200 |

#### 2.3.2 Fine-Tuning Pretrained Models to Enhance Disease Similarity Measurement

For the model to better handle the disease semantic similarity computation task while maintaining its original performance, we utilized MeSH descriptors from category C (diseases) to create a dataset containing pairs of diseases and their disease similarity. The distribution of disease similarities in the dataset we created closely mirrors the disease semantic similarity distribution produced by the ‘multi-qa-MiniLM-L6-cos-v1’ model without fine-tuning. Both types of distributions follow a log-normal distribution pattern. The methods are as follows:

- Determine the depth of the disease descriptors in the MeSH Tree:

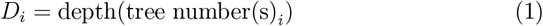
- Determine the depth of the lowest common ancestor (LCA) between two disease descriptors:

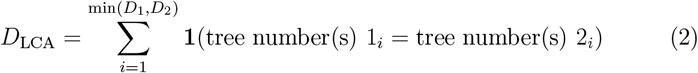

##### Parameter Definitions

tree number(s)1_*i*_ = tree number(s)2_*i*_ indicates the level at which the two diseases share a common node in their hierarchical structure. *i* ranges from 1 to the depth at which the two codes’ lowest common ancestor (LCA) of the two tree numbers is located.

- Calculate the total depth:

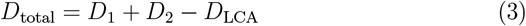
- Calculate the disease similarity:

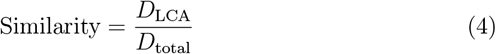
- Calculate the adjusted disease similarity:

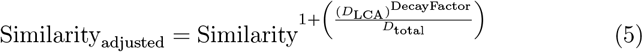

A decay factor of 0.8 was selected in this study. According to the provided formula, the weighted disease similarity between the descriptor: Cardiovascular Infections, tree number(s): C14.260, and the descriptor: Out-of-Hosptial Cardiac Arrest, tree number(s): C14.280.383.610 is 0.14495593273553914 (Figure 3, Figure 4). The weighted disease similarity was further adjusted to better align with the distribution of the output generated by the ‘multi-qa-MiniLM-L6-cos-v1’ model, resulting in a dataset, named ‘MeSHDS’. We utilized this dataset to fine-tune the SBERT model.

**Figure 4.**
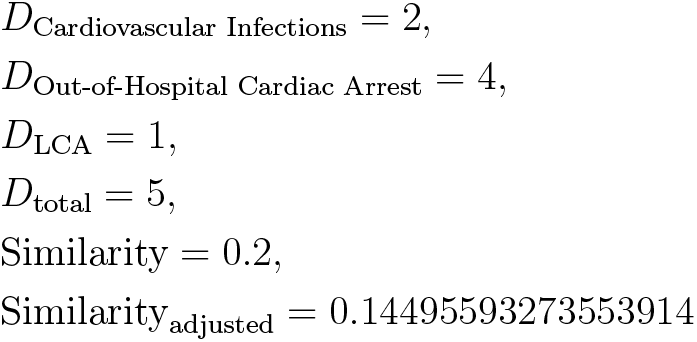
Adjusted Similarity Calculation between ‘Cardiovascular Infections’ and ‘Out-of-Hospital Cardiac Arrest’

#### 2.3.3 Similarity Analysis Between miRNA and Gene

To analyze the similarity between diseases associated with miRNAs and genes, we compute the similarity matrix using two disease string lists: miRNA-associated disease string list and gene-associated disease string list, then analyze the value distribution within the similarity matrix.

##### Semantic String Embedding and Similarity Calculation

First, each string in two lists is embedded using the Sentence-BERT model. The similarity between two strings is obtained by calculating the dot product of their embedding vectors:

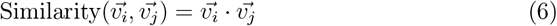

##### Parameter Definitions

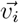 represents the semantic vector of elements *i* in the miRNA-associated disease list embedded with sentence-bert. 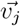 represents the semantic vector of elements *j* in the gene-associated disease list embedded with sentence-bert.

##### Similarity Matrix Calculation

The similarity matrix between miRNA-associated disease and gene-associated disease can be assessed using the following approach:

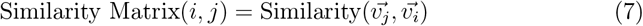

##### Parameter Definitions

Similarity Matrix(*i, j*) represents the element at the *i*-th row and *j*-th column of the similarity matrix.

##### Analyze the Value Distribution in the Similarity Matrix

The similarity matrix of varying dimensions are transformed into a regularized set of length 5 using statistical method, wherein each position represents the distribution of similarities within a specific interval. For example, the first position corresponds to values ranging from 0 to 0.2. This regularized set serves as a classification feature. Meanwhile, by extracting values above 0.8 in the similarity matrix, detailed information on miRNA-associated and gene-associated diseases can be obtained.

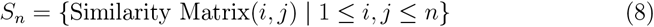

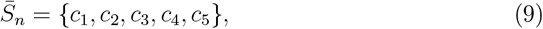

where

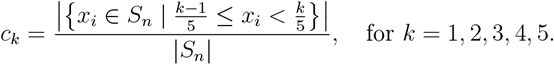

##### Parameter Definitions

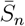 represents the regularized set containing the frequency of similarity values in *S*_*n*_ that fall into each of the 5 intervals. *c*_*k*_ denotes the frequency of similarity values in the *k*-th interval 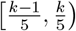.

#### 2.3.4 Identifying Optimal Fine-tuned Models and Classification Methods for Superior Classification Performance

Functional MTIs and MTIs from miRWalk and miRDB were classified based on the regularized set, with the classifier’s performance evaluated using the F1 score. The F1 score, calculated as the harmonic mean of precision and recall, was affected by different fine-tuned models and the choice of classification methods in this study. To ensure optimal performance, we conducted extensive tuning to identify the most suitable model and classification approach. The test set includes 20 genes that are associated with cancer and have sufficient disease data in DisGeNET, namely TP53, PTEN, KRAS, BCL2, FOXO3, KIF2C, PLK1, CENPA, STAT3, MYC, TNF, IL6, TGFB1, IL1B, IL10, IL17A, MMP9, CRP, TLR4, NLRP3 [24–43]. We hypothesize that protein-level experimentally validate MTIs exhibit different disease similarity patterns from predicted MTIs, thus enabling their classification using machine learning techniques. Therefore, we employed five machine learning algorithms, Support Vector Machine, Logistic Regression, Decision Tree, Random Forest, and Gradient Boosting Machine, to classify an equal number of Functional MTIs from miRTarBase and Predicted MTIs from miRWalk & miRDB.

## 3 Results

### 3.1 Performance of Fine-Tuned SBERT Model for Disease Similarity Calculation

The “MeSHDS” dataset comprises 2,249,975 disease pairs along with their corresponding similarities (Figure 5), a substantial volume of data that ensures the high quality of fine-tuning. To test the performance of the fine-tuned SBERT model, we calculated Pearson’s correlation coefficient between the model output scores and scores from BIOSSES and MeSHDS (Figure 6). The outcome demonstrates its capacity to compute biomedical semantic similarity while learning to compute disease similarity through fine-tuning.

**Figure 5.**
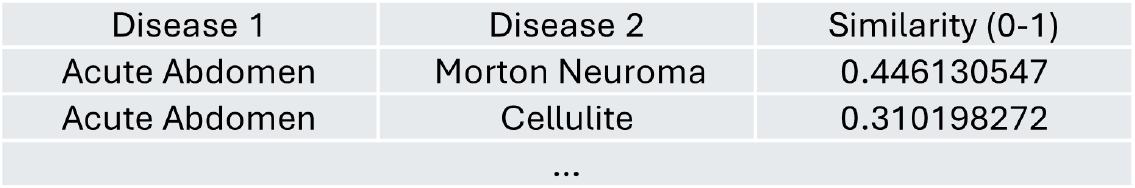
Example of MeSHDS Dataset. Disease 1 and Disease 2 are derived from the MeSH Tree, and their disease similarity is computed using the algorithm described in method.

**Figure 6.**
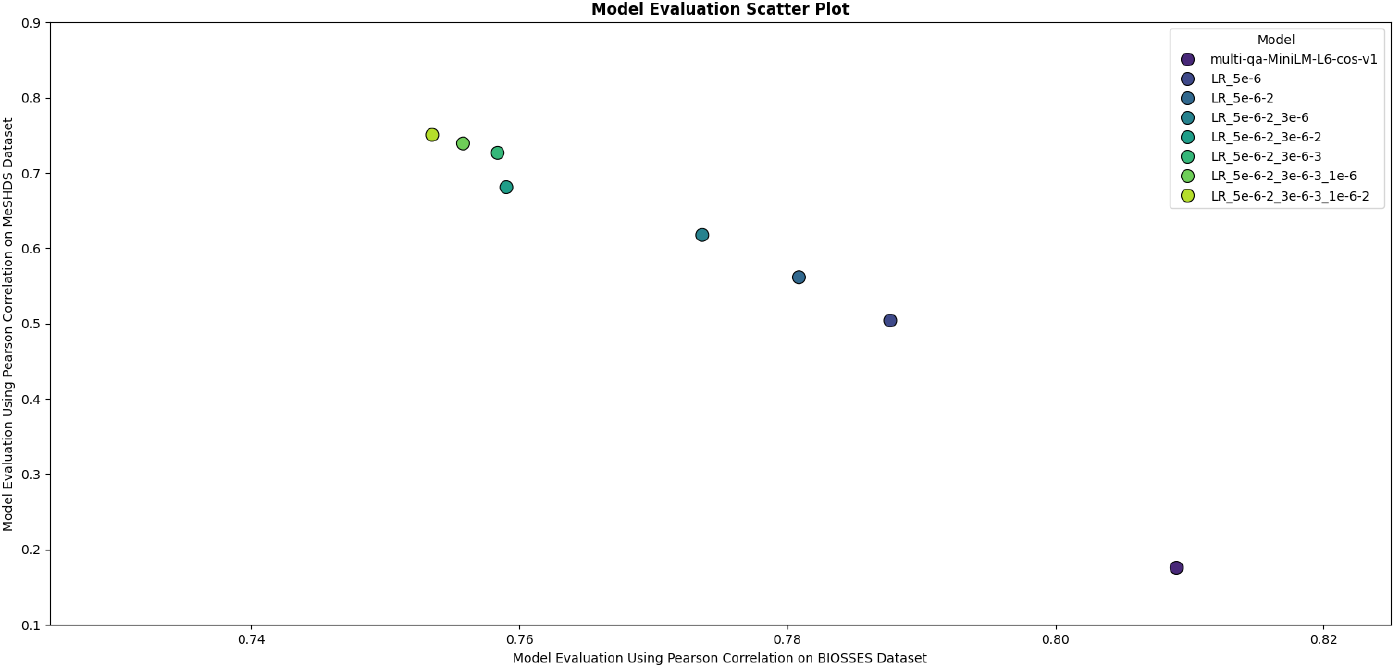
Model Performance Benchmark. The x-axis represents the correlation coefficient between the model-generated similarity scores and the BIOSSES dataset similarity scores, measuring the model’s ability to handle semantic similarity in the biomedical domain. The y-axis represents the correlation coefficient between the model-generated similarity scores and the MeSHDS dataset similarity scores, evaluating the fine-tuning effectiveness. The strategies of fine-tuning are described below:

1. **LR_5e_6**: Fine-tuned for 1 epoch using a learning rate of 5e-6.
2. **LR_5e_6-2**: Fine-tuned for 2 epochs using a learning rate of 5e-6.
3. **LR_5e_6-2_3e_6**: Fine-tuned for 2 epochs with a learning rate of 5e-6, followed by 1 epoch with 3e-6.
4. **LR_5e_6-2_3e_6-2**: Fine-tuned for 2 epochs with a learning rate of 5e-6, followed by 2 epochs with 3e-6.
5. **LR_5e_6-2_3e_6-3**: Fine-tuned for 2 epochs with a learning rate of 5e-6, followed by 3 epochs with 3e-6.
6. **LR_5e_6-2_3e_6-3_1e_6**: Fine-tuned for 2 epochs with a learning rate of 5e-6, 3 epochs with 3e-6, and 1 epoch with 1e-6.
7. **LR_5e_6-2_3e_6-3_1e_6-2**: Fine-tuned for 2 epochs with a learning rate of 5e-6, 3 epochs with 3e-6, and 2 epochs with 1e-6.

### 3.2 Fine-Tuned Model and Classification Method Selection

The F1 score was employed as the primary selection criterion to identify the optimal SBERT model and machine learning method for classifying protein-level experimentally validated MTIs and predicted MTIs. The pre-trained model finetuned using MeSHDS for two rounds, with a learning rate of ‘5e-6’, followed by an additional two rounds of fine-tuning at a learning rate of ‘3e-6’ demonstrated remarkable performance improvement, achieving an 9% increase compared to the original model. The utilization of random forest and gradient descent classification methods in this model leads to optimal overall performance, as evidenced by an F1 Score exceeding 0.8 (Figure 7). After comparison, we have decided to select the ‘LR_5e_6-2_3e_6-2’ model and employ the random forest for further investigation.

**Figure 7.**
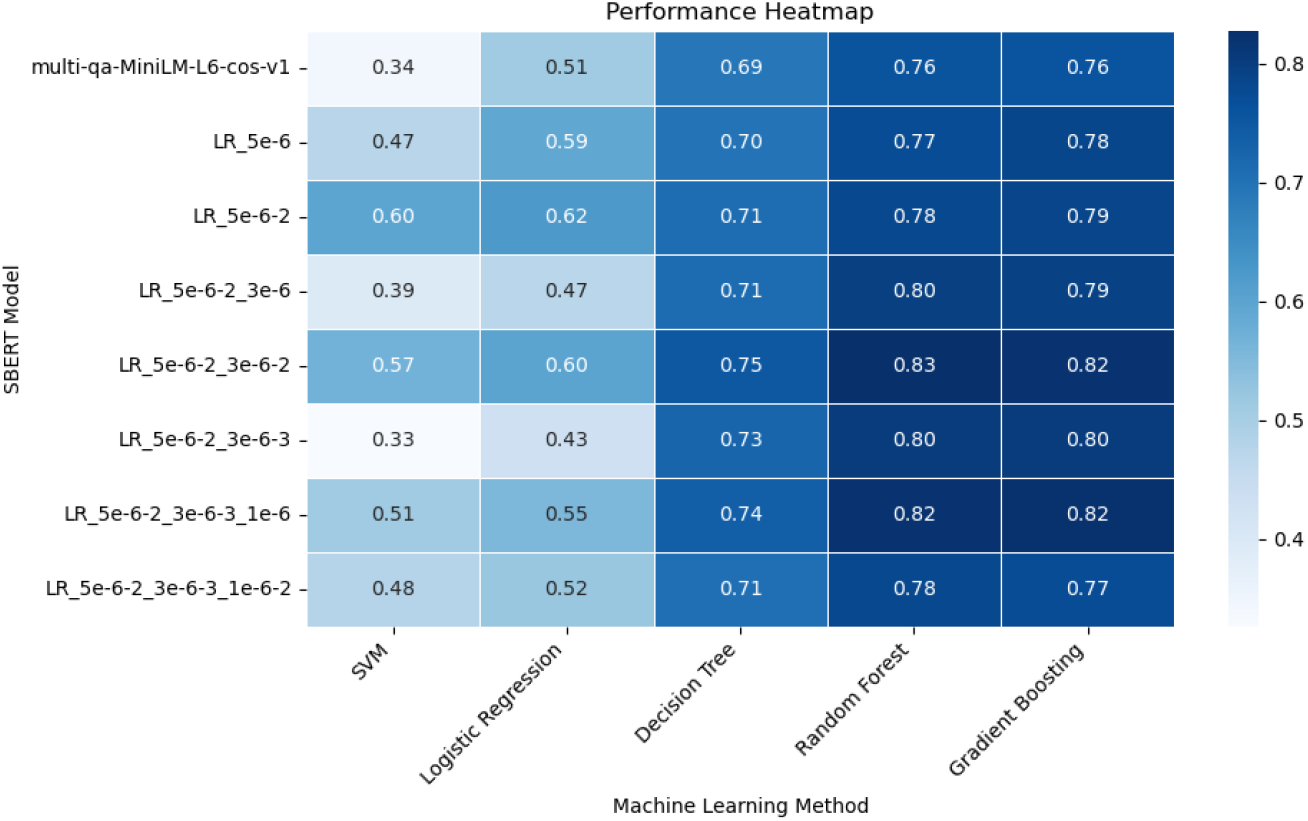
F1 Scores Across Different SBERT Models and Classification Methods. We tested 8 different models using 5 machine learning methods, including Support Vector Machine (SVM), Logistic Regression, Decision Tree, Random Forest, and Gradient Boosting.

### 3.3 MicroRNA and Target Gene Disease Similarity Calculation

The disease similarity matrix of 1,220,904 MTIs was computed, including 1,261 miRNAs from HMDD and 6,085 genes from DisGeNET. The protein-level experimentally validated MTIs from miRTarBase and predicted MTIs from miRWalk and miRDB, were binary classified using a Random Forest algorithm, with the similarity matrix employed as the feature. Random sampling was employed to address the imbalance between functional MTIs and predicted MTIs. The classifier achieved an F1 score of 0.88 (Figure 10, Table 2), with an ROC AUC of 0.94 (Figure 8) and a PRC AUC of 0.93 (Figure 9), demonstrating strong classification performance. Subsequently, the trained Random Forest model was applied to all MTIs, resulting in 1,220,904 predictions. Of these predictions, 189,759 were classified as Protein-level Interaction while 1,031,145 were classified as Prediction Interaction (Figure 11). Furthermore, miRTarDS has accurately identified 90% (518/574) of new functional MTIs in miRTarbase 2025.

**Table 2:** Random Forest Classifier Evaluation Metrics.

| Metric | Formula | Value |
| --- | --- | --- |
| Accuracy | $\frac{TP+TN}{TP+TN+FP+FN}$ | 0.887 |
| Precision | $\frac{TP}{TP+FP}$ | 0.906 |
| Recall | $\frac{TP}{TP+FN}$ | 0.872 |
| F1 Score | $\frac{2 \cdot \text{Precision} \cdot \text{Recall}}{\text{Precision} + \text{Recall}}$ | 0.889 |

**TP:** True Positives (Correctly Predicted Positives)
**TN:** True Negatives (Correctly Predicted Negatives)
**FP:** False Positives (Incorrectly Predicted Positives)
**FN:** False Negatives (Incorrectly Predicted Negatives)

**Figure 8.**
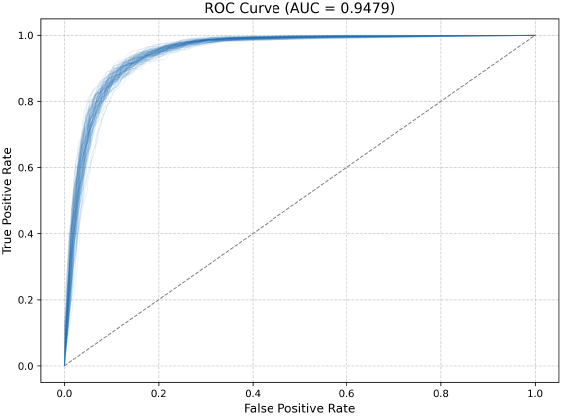
ROC Curve of Random Forest Classifier (100 Repetition)

**Figure 9.**
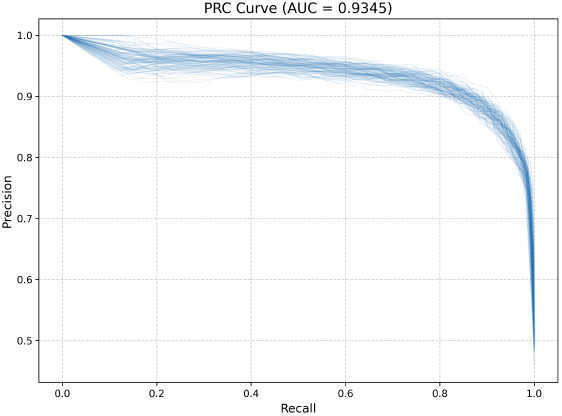
PRC Curve of Random Forest Classifier (100 Repetition)

**Figure 10.**
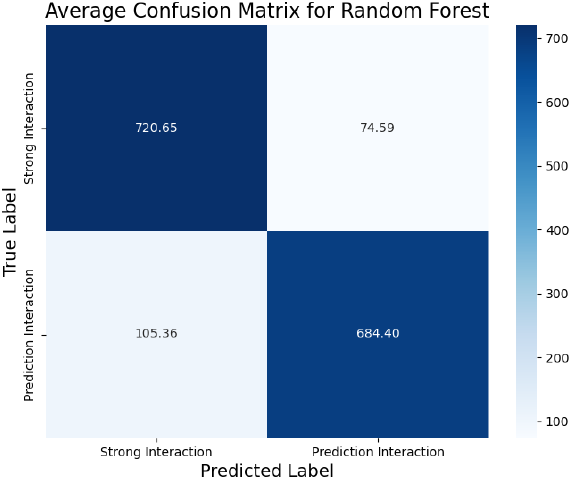
Confusion Matrix Analysis. 4:1 Train-Test Split with 100 Repetition

**Figure 11.**
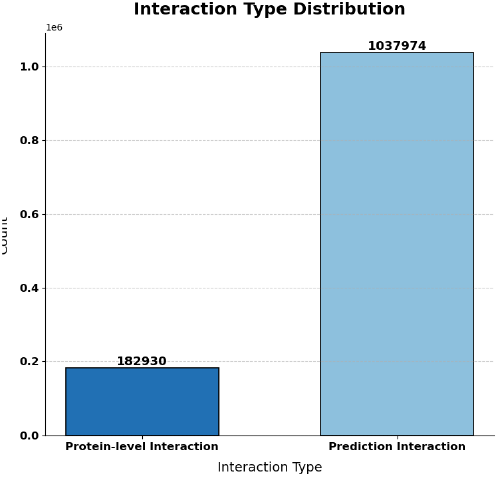
Distribution of Predicted Label

### 3.4 Model Generalizability Across Different Databases

The generalizability of the method was assessed by testing its performance on different databases. We manually downloaded the gene-associated disease data from the KEGG database, and within the 20 previously mentioned genes, only TP53, PTEN, KRAS, MYC, TNF, IL6, IL10, and CRP have enough gene-associated diseases in KEGG. These genes were selected to generate 5,263 MTIs’ disease similarity matrices with both the DisGeNET and the KEGG databases. The correlation coefficients between the regularized set of each MTI across the two databases were calculated. The average Pearson’s correlation coefficient is approximately 0.90 (Figure 12). This result provides evidence that the proposed method exhibits perfect generalizability across different databases.

**Figure 12.**
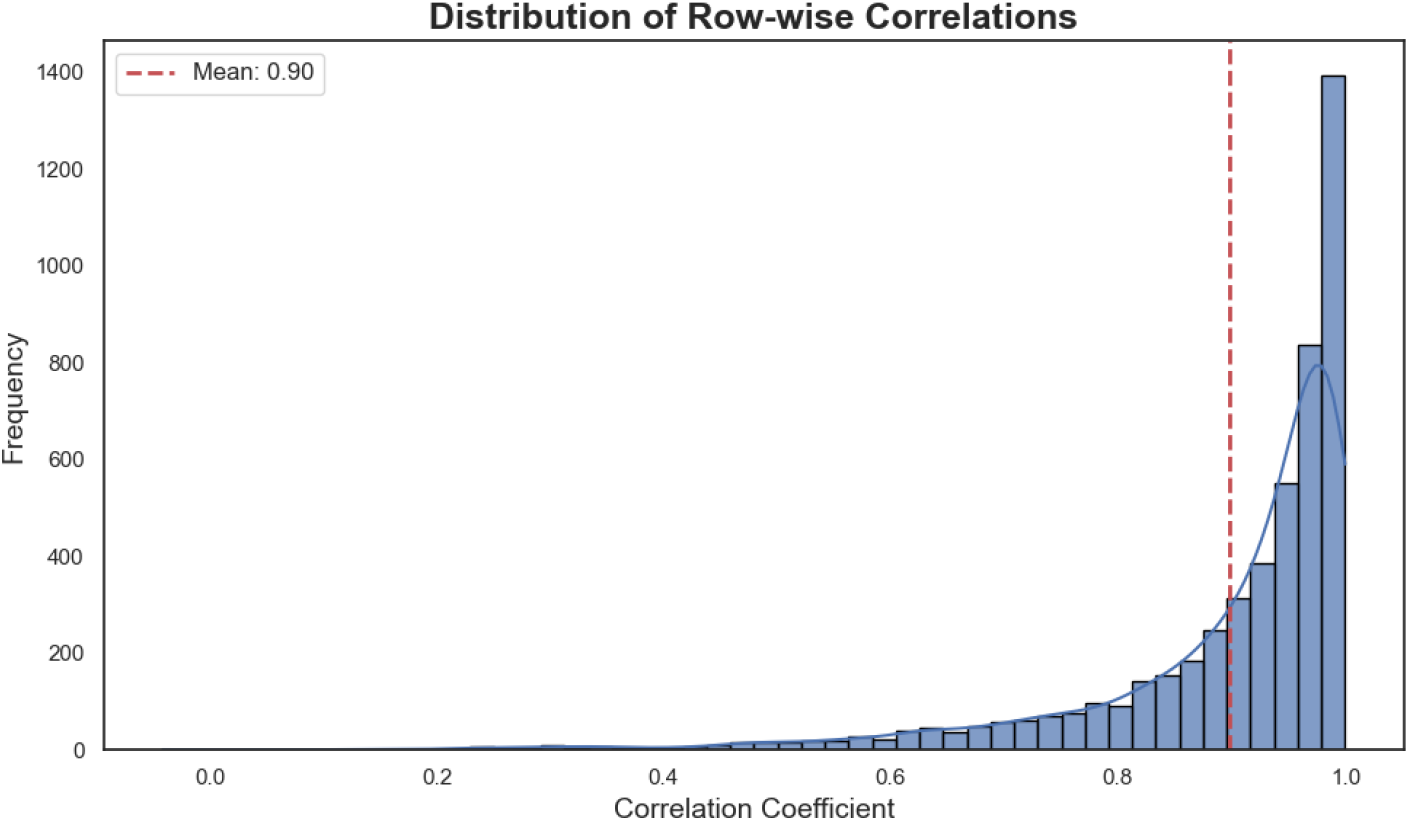
Distribution of Row-wise Correlation Coefficients for Regularized Set Calculated Using DisGeNET and KEGG Databases. This figure compares the disease similarity matrices generated by our method using the same miRNA associated disease database (HMDD) but different gene associated disease databases (DisGeNET and KEGG). More than 5,000 MTIs were calculated. Despite significant differences in the number of gene associated diseases between DisGeNET and KEGG, the correlation coefficients between their results are notably high.

## 4 Discussion

The present study introduces a method to refine sequence-based predicted MTIs by considering the disease similarity between miRNAs and genes. This approach ultimately leads to the development of a database that predicts potential protein-level MTIs and identifies diseases associated with these MTIs. We compared the average disease similarity value distribution of functional MTIs from miRTarbase with predicted MTIs from miRWalk and miRDB (Figure 13), uncovering a slight difference between the two datasets. In the 0–0.2 range, functional MTIs exhibited a lower frequency compared to MTIs from miRWalk and miRDB. However, the trend was reversed in the 0.2–0.4 range, suggesting that MTIs experimentally validated at the protein level tend to show higher similarity between miRNA-associated diseases and gene-associated diseases. This confirms our previous hypothesis.

**Figure 13.**
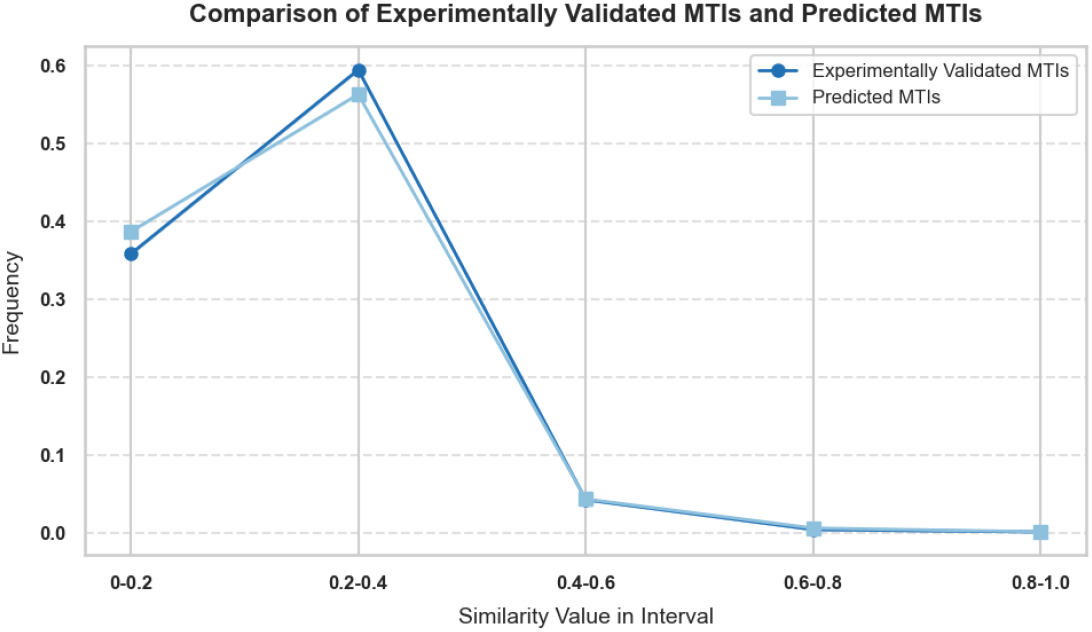
Average Disease Similarity Value Distribution of MTIs from miRTarbase and miRWalk & miRDB. This figure presents the average frequency distribution of similarity values between miRNA-associated diseases and gene-associated diseases for both experimentally validated MTIs and predicted MTIs. Experimentally validated MTIs are confirmed using western blot or reporter assay, while predicted MTIs are derived from computational predictions based on sequence data. The x-axis represents the intervals of similarity values, and the y-axis indicates the average frequency of these similarity values within each interval. The figure demonstrates that diseases associated with miRNAs and genes in experimentally validated MTIs show greater similarity compared to those in predicted MTIs.

Moreover, considering the abundance of miRNA-associated disease and gene-associated disease data, our proposed method holds great potential for widespread application. Notably, the DisGeNET database encompasses 21,671 genes linked to various disease types, while HMDD provides information on 1,900 miRNAs and their associated diseases. The estimated total number of human genes is approximately 22,333 [44], while the estimated number of mature miRNAs in humans is around 2,300 [45]. Based on this estimation, the method can encompass a substantial proportion of both miRNAs and genes. More importantly, compared to the previous disease similarity algorithms, such as MISIM [46] and SemFunSim [47], the SBERT-based disease similarity computation method does not require the input must be present on the MeSH Tree, which greatly improves the scope of computation.

The results of the experiment and the model’s performance on BIOSSES and MeSHDS datasets demonstrate that the Natural Language Processing (NLP) method excels in calculating semantic string similarity in biomedical contexts after specific fine-tuning. The method demonstrates promising scalability and can also be applied to compute disease similarity or functional similarity between other biomolecules, thereby facilitating the exploration of their interactions. In addition to predicting interactions based on physical and chemical properties, the method provides novel insights into disease semantic similarity and functional semantic similarity, thereby optimizing the analysis.

## Declarations

## Acknowledgments

I would like to express my heartfelt gratitude to Dr. SH Xiang. I deeply appreciate the unwavering support from Shenzhen X-institute. I also acknowledge Dr. HY Huang for his feedback during the very early stage. Please note that the data presented in this study may not be integrated into any databases without prior consent from the author.

## Availability of Data and Materials

The MeSHDS dataset and optimal fine-tuned model can be obtained from `Baiming123/MeSHDS` and `Baiming123/Calcu_Disease_Similarity` on HuggingFace, respectively. miRTarDS source data are available at `AltasChen/miRTarDS` on GitHub.

